# M-Ionic: Prediction of metal ion binding sites from sequence using residue embeddings

**DOI:** 10.1101/2023.04.06.535847

**Authors:** Aditi Shenoy, Yogesh Kalakoti, Durai Sundar, Arne Elofsson

**Affiliations:** Science for Life Laboratory and Department of Biochemistry and Biophysics, Stockholm University, 171 21 Solna, Sweden; Department of Biochemical Engineering & Biotechnology, Indian Institute of Technology (IIT) Delhi, New Delhi 110016, India; Yardi School of Artificial Intelligence, Indian Institute of Technology (IIT) Delhi, New Delhi 110016, India

**Author notes:** The authors wish it to be known that the first two authors should be regarded as Joint First Authors.

## Abstract

**Motivation:** Understanding metal-protein interaction can provide structural and functional insights into cellular processes. As the number of protein sequences increases, developing fast yet precise computational approaches to predict and annotate metal binding sites becomes imperative. Quick and resource-efficient pre-trained protein language model (PLM) embeddings have successfully predicted binding sites from protein sequences despite not using structural or evolutionary features (multiple sequence alignments). Using residue-level embeddings from the PLMs, we have developed a sequence-based method (M-Ionic) to identify metal-binding proteins and predict residues involved in metal-binding.

**Results:** On independent validation of recent proteins, M-Ionic reports an area under the curve (AUROC) of 0.83 (recall=84.6%) in distinguishing metal-binding from non-binding proteins compared to AUROC of 0.74 (recall =61.8%) of the next best method. In addition to comparable performance to the state-of-the-art method for identifying metal-binding residues (Ca^2+^, Mg^2+^, Mn^2+^, Zn^2+^), M-Ionic provides binding probabilities for six additional ions (i.e., Cu^2+^, Po_4_^3-^, So_4_^2-^, Fe^2+^, Fe^3+^, Co^2+^). We show that the PLM embedding of a single residue contains sufficient information about its neighbours to predict its binding properties.

**Availability and Implementation:** M-Ionic can be used on your protein of interest using a Google Colab Notebook (https://bit.ly/40FrRbK). GitHub repository (https://github.com/TeamSundar/m-ionic) contains all code and data.

**Supplementary information:** Supplementary data are available at *Bioinformatics* online.

## 1. Introduction

Proteins containing metal cofactors (metalloproteins) are involved in various cellular biochemical processes, including protein folding (Palm-Espling et al., 2012), enzymatic reactions (Andreini et al., 2009), and signalling (Maret, 2017). Understanding metal-binding specificity can provide information about the functions and mechanisms of proteins. Of all proteins, about one-third of them are known to interact with metal ions (Tainer et al., 1991; Andreini et al., 2009). However, as per (Aptekmann et al., 2022), only ∼14% is annotated as metal binding in the UniProt database (The UniProt Consortium, 2017). The size of metal ions, protein activation by a variety of ions (metal promiscuity (Eom and Song, 2019)), and protein precipitation while dealing with high concentrations of samples (Witkowska and Rowińska-Żyrek, 2019) are some of the difficulties involved with experimental methods making them labour-intensive and time-consuming.

Multiple attempts to develop in-silico alternatives for identifying metal-ion binding sites have been previously proposed. These prediction methods can be broadly categorised into three categories, namely (i) sequence-based, (ii) structure-based, and (iii) both sequence/structure-based based on the features used for training. Structure-based methods such as MIB2 (Lu et al., 2022) consider the 3-dimensional neighbourhood surrounding the metal ion. While sequence-based methods like LMetalSite (Yuan, Chen, Wang, et al., 2022), mebi-pred (Aptekmann et al., 2022), and IonSeq (Hu et al., 2016) make use of the protein sequence for training the estimator, IonCon (Hu et al., 2016) employs both sequence and structural information. However, these methods are limited by the availability of template structures and coverage of essential metal ions. Embeddings extracted from various protein language models (PLMs), such as ESM-1b (Rives et al., 2021) and ProtTrans (Elnaggar et al., 2022), are quick to generate and have shown good performance for identifying subcellular localisation (Stärk et al., 2021), post-translational modification (Pokharel et al., 2022) and ligand binding sites (Littmann et al., 2021).

In this study, residue-level PLM embeddings were employed to estimate (i) the likelihood of a given metal ion binding to a protein (protein level prediction) and (ii) the probability of individual protein residues interacting with the given metal ion (residue level prediction). M-ionic provides binding-site predictions for the ten most frequently occurring metal ion groups with residue-level binding probabilities for multiple ions simultaneously. Despite using only single residue embedding, M-Ionic performs comparably to recent metal-binding predictors such as mebi-pred (Aptekmann et al., 2022) and LMetalSite (Yuan, Chen, Wang, *et al*., 2022). Therefore, we show that residue embeddings can effectively predict metal-ion binding sites.

## 2. Materials and methods

### 2.1. Datasets

In this study, we have used three sets of metal-binding proteins. As in earlier studies (Hu et al., 2016; Cao et al., 2017; Yuan, Chen, Wang, et al., 2022), proteins from BioLip were used for training. Secondly, two independent sets were used for validation, one consisting of newer proteins in BioLip and the benchmark set from MIonSite. Finally, we also created a negative set from PDB. These are all described in detail below.

#### 2.1.1. Training, validation, and independent test set

Metal binding proteins (n=29404) were curated from the BioLip database (Yang *et al*., 2013) on 2022-09-27. Proteins having sequence length >1000, resolution (>3Å), and ions with less than 1000 protein entries were removed. (Table S1). Sequences were clustered using CD-HIT (Li and Godzik, 2006) at 100% sequence identity to remove identical sequences with contradictory labels. Previous studies (Krissinel, 2007; Pearson, 2013) have shown that homology exists between proteins having 30% sequence identity. Hence, we performed homology reduction using a sequence similarity threshold of 20% using BlastClust (Altschul *et al*., 1990) with the parameters -p T -L .9 -b T -S 20. The 6203 clusters obtained were shuffled and split into six partitions. One partition (TestFold6) (n=3002) was reserved upfront as the independent test set. The remaining five partitions (n=26407) were used to train the estimator using a five-fold cross-validation strategy. Note: TestFold6 was not used during the training and validation of M-Ionic and had no homology with the training datasets. M-Ionic performance was further benchmarked on two independent datasets: (1) Recent BioLip (n=5377) consisting of only recent proteins and (2) MIonSite Benchmark dataset (n=267) (Qiao and Xie, 2019).

#### 2.1.2. Recent BioLip proteins

Proteins from the BioLip dataset (Yang *et al*., 2013) after 2022-09-27 till 2023-05-31 were used as an independent test set. Metal ions having a minimum of 1000 proteins and each having resolution (<3Å) were selected. Identical sequences (100% sequence identity threshold) were removed using CD-HIT (Li and Godzik, 2006). Since these proteins are outside the training/testing datasets of M-Ionic, mebi-pred (Aptekmann *et al*., 2022), and LMetalSite (Yuan, Chen, Wang, *et al*., 2022), the ‘Recent BioLip’ dataset (n=5377) allows for unbiased testing.

#### 2.1.3. Independent benchmark dataset from MIonSite

The performance of M-Ionic was benchmarked against other metal-binding predictors (LMetalSite (Yuan, Chen, Wang, et al., 2022); GASS-Metal (Paiva et al., 2022), IonSeq and IonCon (Hu *et al*., 2016); TargetS (Yu et al., 2013); MetalDetector (Lippi et al., 2008)) using the independent validation dataset from MIonSite (Qiao and Xie, 2019). Note: The benchmark testing dataset from MIonSite (n=267) differs from the M-Ionic testing set (TestFold6) (n=3002) described above. The two sets have an overlap of only 10 PDB chains (5y9e_E, 5z68_B, 5z9x_A, 6c1j_A, 6c33_A, 6ci7_B, 6cko_C, 6d0y_C, 6ekb_A, 6eke_B).

#### 2.1.4 Negative metal binding dataset

To evaluate whether the method can distinguish between (metal) binding and non-binding proteins, a negative set (i.e., protein chains that do not bind to any metal ion) was created. Protein bio-assemblies from Protein Data Bank (PDB) (Berman et al., 2000) with less than six protein chains and chains with no metals in the HETATM records were filtered (n=3810) and clustered at 100% identity. The resulting dataset (n=3224) was used alongside the ‘Recent BioLip’ dataset (Table 1; Figure 2a-b) and TestFold6 (Table S2; Figure S1a-b) to plot the ROC curves.

**Table 1.** Performance on distinguishing metal binding and non-binding proteins using the ‘Recent BioLip’ dataset and the negative set.

| Methods | Prec. | Rec. | Acc. | F1 | MCC | AUPR | AUROC |
| --- | --- | --- | --- | --- | --- | --- | --- |
| Mebi-pred (2022) | **0.39** | 0.56 | **0.84** | 0.46 | 0.38 | 0.27 | 0.72 |
| LMetalSite (2022) | 0.38 | 0.62 | 0.83 | 0.47 | 0.39 | 0.28 | 0.74 |
| M-Ionic (Current) | 0.39 | **0.85** | 0.82 | **0.53** | **0.49** | **0.35** | **0.83** |

**Figure. 1.**
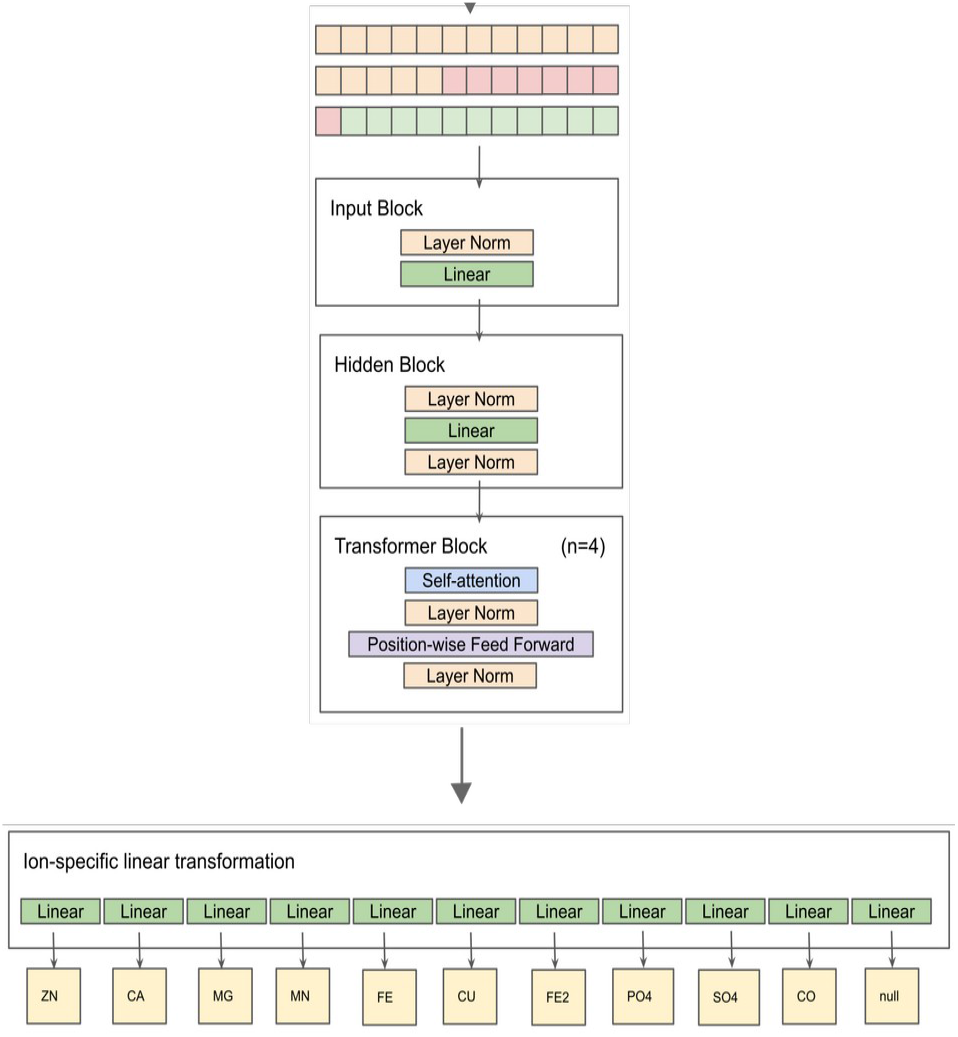
The workflow of M-Ionic consists of an input block, a hidden block and four transformer blocks. Residues are batched together and trained along the residue-embedding dimension. The final ion-specific linear layers provide output probabilities of multiple ions for each residue.

**Figure. 2.**
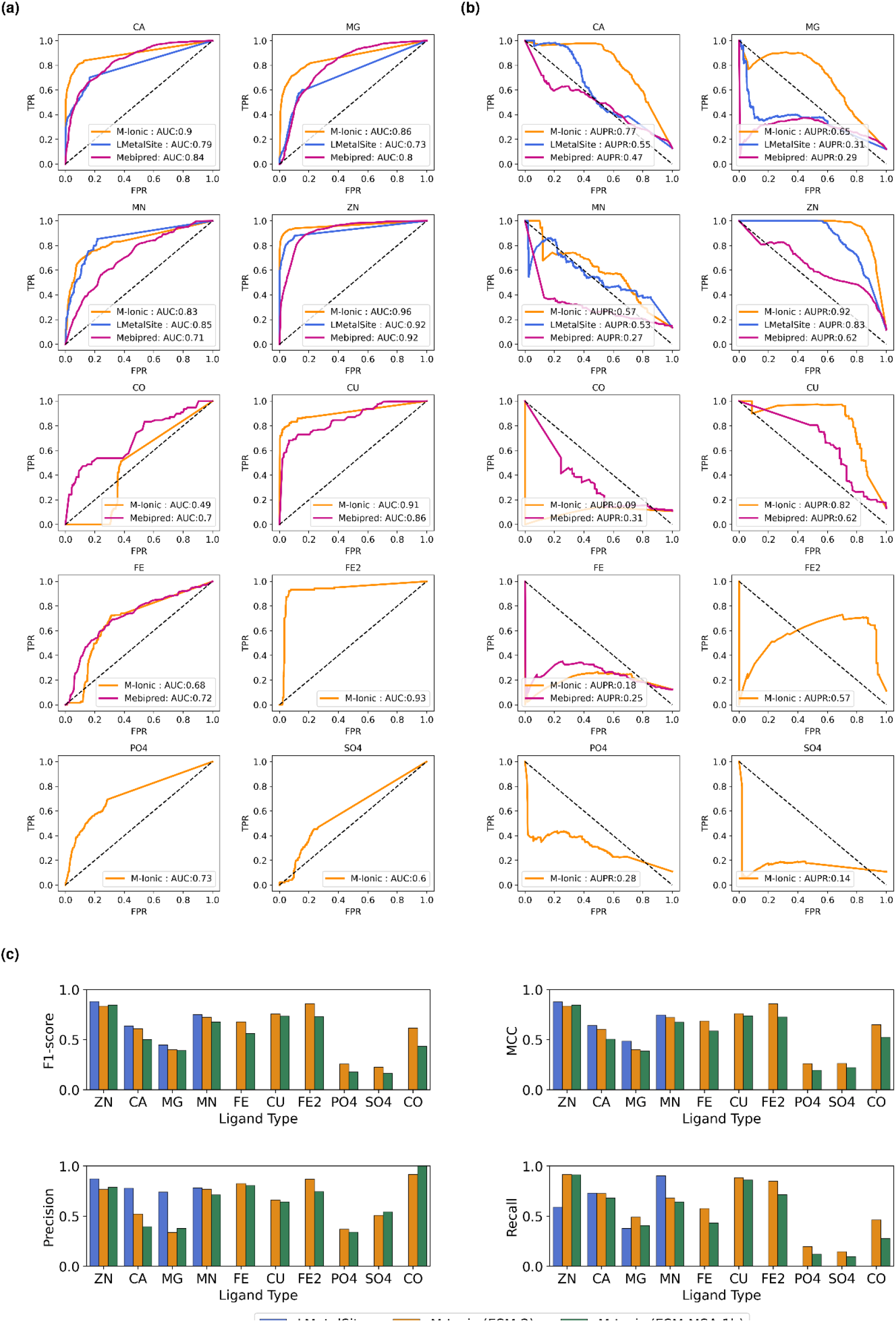
(a) Protein-level: Comparison of ROC curves for the performance of each ion type for M-Ionic (this study) trained on ESM-2 embeddings, LMetalSite (Yuan, Chen, Wang, et al., 2022) and mebi-pred (Aptekmann et al., 2022) (b) Protein-level: Comparison of Precision-Recall curves for the performance of each ion type for each of the methods (c) Residue-level: MCC scores of performance of M-Ionic (trained on ESM-2 and ESM-MSA-1b embeddings) and LMetalSite (Yuan, Chen, Wang, et al., 2022) on ‘Recent BioLip’ dataset (i.e., outside training data).

### 2.2. Feature extraction

#### 2.2.1. Sequence representation

ESM-2 (Rives et al., 2021) and ProtT5 (Elnaggar et al., 2022) residue-level embeddings were generated for all sequences. The dimension of each ESM-2 (esm2_t33_650M_UR50D) protein embedding was *Lx1280*, and that of ProtT5 (ProtT5-XL-UniRef50) protein embedding was *Lx1024*, where L is the length of the protein (i.e. each residue has a 1280 or 1024 dimension embedding).

#ESM-2

python3 esm/scripts/extract.py

esm2_t33_650M_UR50D input.fasta esm_embeddings

--repr_layers 33 --include per_tok #ProtT5

python3 Embedding/prott5_embedder.py --input input.fasta --output proT5_embeddings.h5

#### 2.2.2. MSA representation

Multiple sequence alignments (MSA) were generated using HHblits (Remmert et al., 2012) version 3.3.0 with three iterations against UniClust30 (Mirdita et al., 2017) (version Uniclust30_2018_08).

hhblits -i äFASTA -oa3m äOUTPUT -n 3 -d uniclust30_2018_08/uniclust30_2018_08 -cpu 32

The MSA was converted into a one-hot MSA and reweighted with a cutoff=0.8 to generate the position-specific scoring matrices (PSSM) (Adhikari, 2020). The MSAs were also converted to ESM-MSA-1b (esm_msa1b_t12_100M_UR50S) protein embedding, i.e., *Lx768* dimensional vector where L is the protein sequence length.

#### 2.2.3. Structural properties

An *Lx14* feature DSSP vector was generated as in (Yuan, Chen, Rao, *et al*., 2022). This vector has three components: (a) One-hot encoding for eight secondary structure states along with one dimension for unknown states (*Lx9*), (b) sine and cosine transformations of the torsion backbone phi/psi angles (*Lx4*), and (c) relative solvent accessibility (*Lx1*), where L is the length of the protein.

### 2.3. Implementation

Each embedding was fed as input to the network one residue at a time. The M-Ionic architecture consists of four blocks: input, hidden, encoder, and ion-specific blocks. It starts with an input block that takes the protein features as input and applies layer normalisation, followed by a linear transformation with ReLU as an activation function. The hidden block processes the features by applying layer normalisation, dropout regularisation, and another linear transformation. The encoder layer comprises four transformer modules consisting of self-attention and position-wise feed-forward layers. Lastly, a linear transformation for each ion (10) and one exclusive dimension (1) for designating the non-binding status of the residue make up the ion-specific block. These linear layers consist of linear transformations followed by leaky ReLU activation functions. The complete model architecture is available on the associated GitHub repository and depicted graphically in Figure 1. In summary, the model utilised self-attention (4 attention heads) and feed-forward networks within the transformer architecture to effectively process protein features and make accurate predictions for multiple ions. The model was trained for 30 epochs using the Adam optimiser and BCEWithLogits Loss with a learning rate of 0.001. The output is an *Lx11* dimensional vector with the binding probabilities of multiple metal ions for each residue in a protein of length L.

### 2.4. Evaluation metrics

Precision, recall, F1-score, and Matthew’s correlation coefficient (MCC) were used to evaluate how methods perform on metal-binding site prediction.

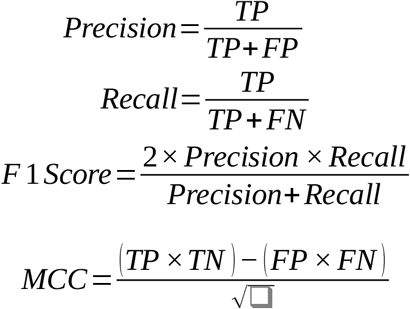

TP represents the number of true positives (metal-binding residues predicted correctly), FP represents the number of false positives (non-metal-binding residues predicted as metal binding), TN represent the number of true negatives (non-metal-binding residues predicted correctly), and FN represents the number of false negatives (metal-binding residues predicted as non-metal-binding). Additionally, to evaluate protein-level prediction (ability to distinguish between metal-binding and non-binding proteins), we used receiver operating characteristic (ROC) and reported the area under the ROC curve (AUROC) score. Precision-recall (PR) curves and area under the PR curves (AUPR) were also reported.

## 3. Results and discussion

The evaluation of the metal-binding prediction methods was conducted at two levels: (1) protein level (i.e., whether the method could predict if the protein had a likelihood to bind to metals or not) (2) residue-level (i.e., the ability to predict the probability of each residue binding to multiple metal ions).

### 3.1. M-Ionic identifies metal-binding proteins

Receiver operating characteristic (ROC) curves and precision-recall (PR) curves were employed to evaluate whether methods could correctly distinguish proteins that bind to metals and those that do not. M-Ionic produces the binding probabilities for each of the ten metal ions for each residue in the protein. In contrast, mebi-pred (Aptekmann et al., 2022) generated only protein-level predictions (i.e., the probability of any protein sequence binding to a metal ion). LMetalSite (Yuan, Chen, Wang, et al., 2022), like M-Ionic, gave binding probabilities at the residue level for four ions (Ca^2+^, Mg^2+^, Mn^2+^, Zn^2+^) as well as 0 or 1 logits indicating the metal-binding status of each residue. The corresponding predicted probabilities of the residues indicated with logit=1 were used in the ROC curve. LMetalSite and M-Ionic differ in their inputs and how the network is trained. LMetalSite batches proteins of different lengths, pads them together, and trains along the length of the proteins. M-Ionic, however, is trained on the embedding dimension (i.e. 1280 for ESM-2). Figure 2a-b shows the performance of M-Ionic compared against recent methods mebi-pred (Aptekmann et al., 2022) and LMetalSite (Yuan, Chen, Wang, et al., 2022) on the ‘Recent BioLip’ (and the negative dataset). We see that M-Ionic reports a higher true positive rate (TPR) at false positive rates (FPR) (except for Co and Fe) compared to other methods and a 30% increase in MCC (Table 1). A similar performance was observed when tested on the independent test set -TestFold6 (and the negative dataset) (Table S2; Figure S1a-b).

### 3.2. M-Ionic identifies metal-binding residues

M-Ionic identifies residues interacting with multiple ions (residue-level prediction) at comparable performance with LMetalSite based on F1-score and MCC (Figure 2c). M-Ionic has better recall (especially for Zn^2+^ and Mg^2+^) but has lower precision values than LMetalSite. Table S3 shows the performance of M-Ionic (trained with ESM-2) with other evaluation metrics. The results from the benchmark set from MIonSite (Qiao and Xie, 2019) are reported in Table S4, showing that for the eight metal ions compared, M-Ionic performs comparably to the other predictors. A similar performance was observed when the methods were tested on the M-Ionic independent test set -TestFold6 (Table S5 and Figure S1c).

### 3.3. Ablation experiments

Three classes of features were used to train the network: (a) Sequence-based (ESM-2, ProtT5), (b) MSA-based (PSSM, ESM-MSA-1b), and (c) Structural properties (DSSP). We did not see an improvement in performance using ProtT5 residue-level embeddings (Table S5).

#### 3.3.1. Impact of evolutionary information

We observed that certain metal ions, e.g., iron and zinc, have a higher binding affinity to certain amino acids (e.g. cysteine, histidine) (Figure S2), as shown in (Cao et al., 2017; Barber-Zucker et al., 2017). However, no improvement was observed using explicit evolutionary information to improve the residue-level prediction performance, i.e., ESM-MSA-1b in Figure 2c. The 22-dimensional PSSM feature vector resulted in a lower performance than the model trained with ESM-2, whereas the 768-dimensional ESM-MSA-1b embedding performed at par with ESM-2. When the MSA-based features were combined with ESM2 features, a slight increase in performance was observed (Table S3).

#### 3.3.2. Impact of structural information

Just as with the performance of the one-hot encoding of the protein sequence, the 14-dimensional DSSP vector yielded null performance when trained with the same architecture. We used combinations of the DSSP features with ESM-2 residue embeddings. However, there was a drop in the performance for feature combinations involving structural features (Table S5). The M-Ionic architecture (Figure 1) relies on the information within each residue embedding. The DSSP vector for an individual residue comprises that residue’s secondary structure and bond angles. Since it does not contain context-dependent neighbourhood information, using structural features on a residue level does not yield better results.

### 3.4. Single residue embeddings can predict metal-binding sites

To verify whether the embeddings from neighbouring residues influenced the prediction of the residue of interest, we retrained the M-Ionic network with batch size 1 (ESM-2 (Batch-1), i.e., the network is trained one-residue at a time) and by scrambling the order of residues within a batch of 128 residues (ESM-2 (Scrambled Batch-128), i.e., scrambling the order of the residues from the original protein sequence). No difference between the models was observed, revealing that the network uses only single residue information (Figure 3 and Table S6). Therefore, M-Ionic shows that a single residue embedding already contains neighbourhood information and is sufficient to predict the metal binding probability for that residue.

**Figure. 3.**
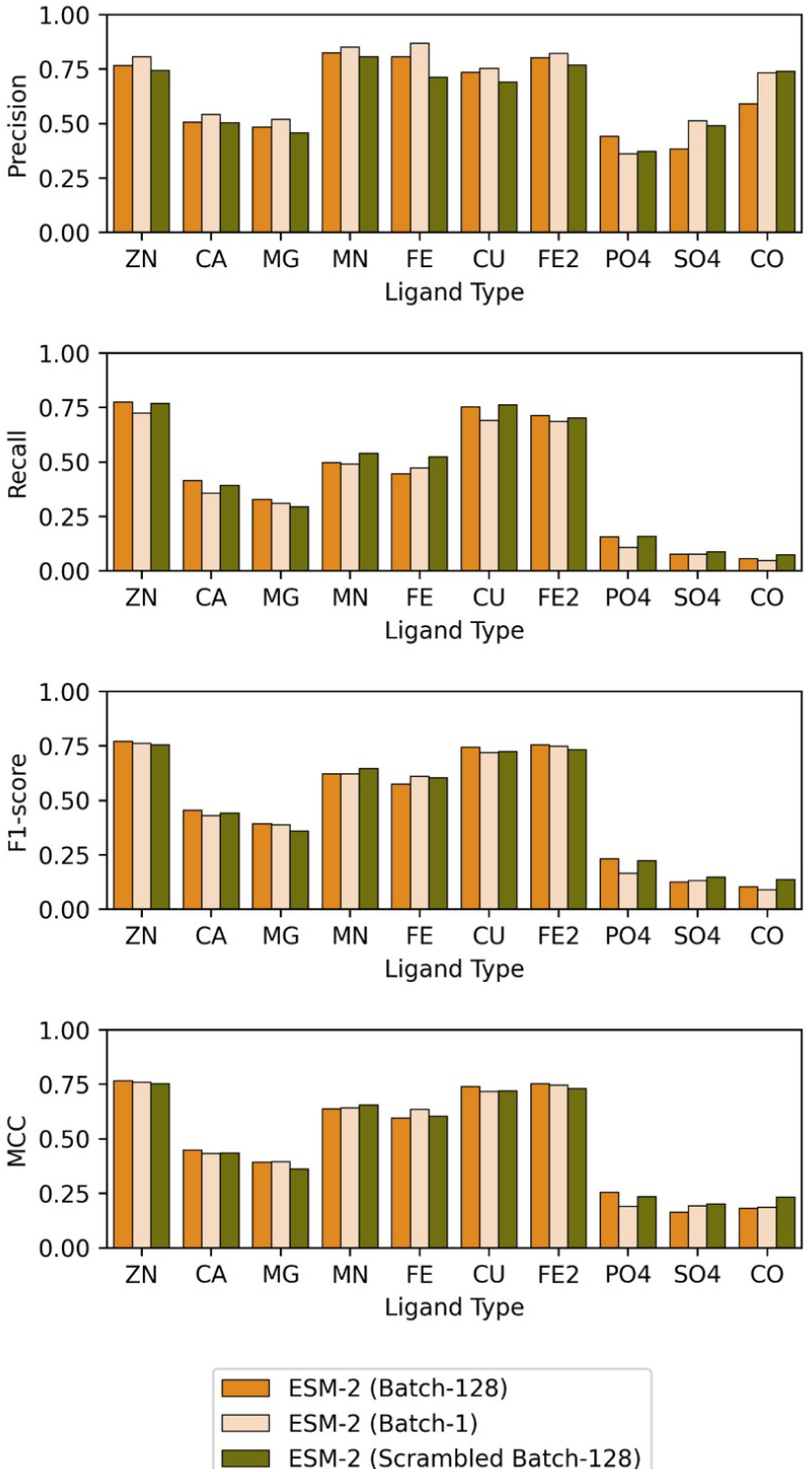
Evaluation metrics (Precision, Recall, F1, MCC) for the various models trained (a) ESM-2 (Batch-128) is trained with ESM-2 model with batch size 128 as in original M-Ionic implementation (b) ESM-2 (Batch-1) is trained with ESM-2 model with batch size 1 (c) ESM-2 (Scrambled Batch-128) trained on ESM-2 model with batch size 128 but with the residues within each batch were shuffled randomly along the length dimension such that the positions of the residues within the sequence were changed, but the embedding of each residue was untouched.

### 3.5 Performance of mutated binding residues

To further validate our method, we mutated all the residues involved in metal binding to another amino acid and studied the impact of these mutations on predicted probabilities from M-Ionic. We found that the number of predicted true positives dropped when metal binding residues were mutated to neutral residues like alanine or glycine (Figure 4). In contrast, the number of false positives remained the same on average. Cysteine and histidine mutation, however, show many true positive residues. However, we expect this because those residues are frequently involved in binding (especially for Zn^2+^ and Fe^2+^) (Figure S2). When comparing the distribution of predicted probabilities of mutated residues, we observe a shift (towards 0.0) from the probabilities of the binding residues of the non-mutated (original) protein sequence (Figure S3).

**Figure. 4.**
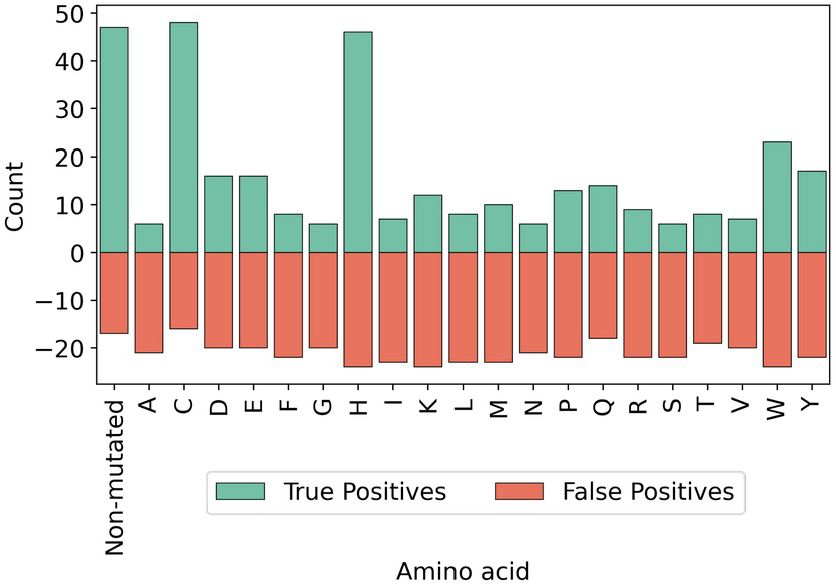
Count plot indicating the number of true and false positives of the M-Ionic predictions. Here, the predictions from the non-mutated sequence are compared against the predictions on the mutated sequence (i.e., when all the binding residues are replaced with one of the amino acids). The False positives remain similar, but when an ion-binding is mutated to anything but a His or Cys virtually, all true binding residues are missed.

### 3.6. Analysis of M-Ionic performance for specific protein subgroups

The performance of M-Ionic was further analysed on specific groups of proteins (DNA-binding; Transmembrane) and according to different taxonomic groups. No differences between performances within the groups were found (p-value =<0.05 obtained using Mann–Whitney T-test) (Table S7; Table S8; Table S9).

## 4. Conclusion

On benchmarking against recent proteins and negative set, the performance of M-Ionic was comparable to mebi-pred at the protein level (i.e., the ability to identify which metals a protein binds to). For the most frequent metal ions (Ca^2+^, Mg^2+^, Mn^2+^, Zn^2+^), M-Ionic reported an MCC score of 0.583 compared to 0.49 (LMetalSite) and 0.42 (mebi-pred), respectively. At the residue level (i.e., the ability to predict which residues are involved in metal-ion-specific binding), M-Ionic generated binding-site predictions (with MCC scores comparable with LMetalSite) for six additional metal ions (i.e. Co^2+^, Cu^2+^, Po_4_ ^3-^, So_4_ ^2-^, Fe^2+^, Fe^3+^). No improvement in performance was observed when ablation experiments with explicit evolutionary and structural features were performed.

This study presents an alternative approach to training pre-trained PLM embeddings by looking within a residue embedding. We show that embeddings from individual residues contain neighbourhood information and have captured long-range dependencies in the pre-training stage. An easy-to-use notebook to generate quick predictions with a probability of binding ten metal ions for each residue in the protein of interest has also been made available.

## Author contributions

AS prepared the datasets. YK developed training architecture and optimised model parameters. AS and YK analysed the results. AS wrote the initial draft of the manuscript. All authors contributed to the manuscript.

## Acknowledgements

We thank Petras Kundrotas and Ho-Yeung Chim for their valuable discussions. We also thank Qianmu Yuan for their input regarding LMetalSite.

## Availability and Implementation

An easy-to-use Google Colab Notebook is freely available at https://bit.ly/40FrRbK. The source code is available from https://github.com/TeamSundar/m-ionic.

