## Supplementary Information for "M-Ionic: Prediction of metal ion binding sites from sequence using residue embeddings"

*To whom correspondence should be addressed.

Figure. S3. Probabilities distributions (output from M-Ionic) showing the effect of mutating metal-binding residues to another residue compared to the non-mutated (original) sequence. 20

**3. References................................................................................................................ 21**

### 1. Tables

**Table S1.** Summary of metal-binding proteins in BioLip dataset

| Ions | Number of protein entries | Number of ions bound with protein receptors | Number of binding residues (P) | Number of non-binding residues (N) | Ratio of binding to non-binding residues (P)/(N) |
|---|---|---|---|---|---|
| $Zn^{2+}$ | 48593 | 1952 | 126278 | 7704568 | 0.016 |
| $Ca^{2+}$ | 38258 | 39147 | 131250 | 6041662 | 0.022 |
| $Mg^{2+}$ | 34958 | 5315 | 78542 | 6959299 | 0.011 |
| $Mn^{2+}$ | 11650 | 2850 | 34315 | 2254077 | 0.015 |
| $Fe^{3+}$ | 6366 | 6038 | 20311 | 1227819 | 0.017 |
| $Cu^{2+}$ | 5617 | 27348 | 16257 | 948823 | 0.017 |
| $Fe^{2+}$ | 3054 | 10444 | 9168 | 601517 | 0.015 |
| $Co^{2+}$ | 2018 | 33127 | 6533 | 369360 | 0.018 |
| $Po_4^{3-}$ | 2759 | 2513 | 12595 | 605193 | 0.021 |
| $So_4^{2-}$ | 2365 | 2291 | 9840 | 432458 | 0.023 |

**Table S2.** Performance on distinguishing metal binding and non-binding proteins using the independent test set generated in this study (TestFold6) and negative set

| Methods | Year | Precision | Recall | Accuracy | F1-score | MCC | AUPR | AUROC |
|---|---|---|---|---|---|---|---|---|
| mebi-pred | *2022* | 0.206 | 0.643 | 0.85 | 0.311 | 0.304 | 0.151 | 0.752 |
| LMetalSite | *2022* | **0.217** | 0.565 | **0.87** | **0.314** | 0.293 | 0.146 | 0.726 |
| M-Ionic | *Current* | 0.18 | **0.855** | 0.787 | 0.297 | **0.329** | **0.161** | **0.819** |

| **Table S3.** Impact of evolutionary (MSA, PSSM) on metal-binding site prediction using 'Recent BioLip' dataset (i.e. independent test set of recent PDB proteins) | | | | | | | |
|---|---|---|---|---|---|---|---|
| Ligand Type | Features | Precision | Recall | F1-score | MCC | AUROC | Average Precision |
| $Zn^{2+}$ | PSSM | 0.587 | 0.818 | 0.684 | 0.685 | 0.902 | 0.485 |
| | ESM-2 + PSSM | 0.759 | 0.917 | 0.831 | 0.830 | 0.955 | 0.698 |
| | ESM-2 + ESM-MSA-1b | 0.791 | 0.924 | 0.852 | 0.851 | 0.959 | 0.733 |
| | ESM-MSA-1b | 0.789 | 0.913 | 0.846 | 0.845 | 0.954 | 0.722 |
| | ESM-2 | 0.766 | 0.915 | 0.834 | 0.833 | 0.954 | 0.703 |
| | LMetalSite | 0.872 | 0.589 | 0.880 | 0.877 | 0.942 | 0.777 |
| $Ca^{2+}$ | PSSM | 0.317 | 0.132 | 0.187 | 0.193 | 0.563 | 0.062 |
| | ESM-2 + PSSM | 0.504 | 0.730 | 0.596 | 0.596 | 0.857 | 0.374 |
| | ESM-2 + ESM-MSA-1b | 0.513 | 0.722 | 0.600 | 0.598 | 0.853 | 0.376 |
| | ESM-MSA-1b | 0.395 | 0.681 | 0.500 | 0.505 | 0.828 | 0.276 |
| | ESM-2 | 0.521 | 0.726 | 0.607 | 0.605 | 0.855 | 0.385 |
| | LMetalSite | 0.779 | 0.730 | 0.637 | 0.642 | 0.768 | 0.429 |
| $Mg^{2+}$ | PSSM | 0.223 | 0.293 | 0.253 | 0.248 | 0.642 | 0.072 |
| | ESM-2 + PSSM | 0.312 | 0.494 | 0.382 | 0.386 | 0.742 | 0.159 |
| | ESM-2 + ESM-MSA-1b | 0.290 | 0.487 | 0.364 | 0.369 | 0.738 | 0.146 |
| | ESM-MSA-1b | 0.377 | 0.406 | 0.391 | 0.385 | 0.700 | 0.158 |
| | ESM-2 | 0.338 | 0.490 | 0.400 | 0.400 | 0.740 | 0.170 |
| | LMetalSite | 0.740 | 0.377 | 0.448 | 0.485 | 0.660 | 0.244 |
| $Mn^{2+}$ | PSSM | 0.534 | 0.262 | 0.351 | 0.369 | 0.630 | 0.148 |
| | ESM-2 + PSSM | 0.742 | 0.692 | 0.716 | 0.714 | 0.845 | 0.517 |
| | ESM-2 + ESM-MSA-1b | 0.778 | 0.649 | 0.708 | 0.708 | 0.823 | 0.509 |

| | | | | | | | |
|---|---|---|---|---|---|---|---|
| | ESM-MSA-1b | 0.713 | 0.642 | 0.676 | 0.673 | 0.820 | 0.462 |
| | ESM-2 | 0.770 | 0.680 | 0.722 | 0.721 | 0.839 | 0.527 |
| | LMetalSite | 0.782 | 0.903 | 0.749 | 0.747 | 0.859 | 0.566 |
| $Fe^{3+}$ | PSSM | 0.783 | 0.176 | 0.288 | 0.368 | 0.588 | 0.148 |
| | ESM-2 + PSSM | 0.785 | 0.589 | 0.673 | 0.677 | 0.793 | 0.468 |
| | ESM-2 + ESM-MSA-1b | 0.832 | 0.538 | 0.654 | 0.666 | 0.769 | 0.454 |
| | ESM-MSA-1b | 0.806 | 0.431 | 0.562 | 0.586 | 0.715 | 0.355 |
| | ESM-2 | 0.825 | 0.575 | 0.678 | 0.686 | 0.787 | 0.480 |
| | LMetalSite | N/A | N/A | N/A | N/A | N/A | N/A |
| $Cu^{2+}$ | PSSM | 0.592 | 0.307 | 0.404 | 0.420 | 0.652 | 0.192 |
| | ESM-2 + PSSM | 0.644 | 0.903 | 0.752 | 0.759 | 0.948 | 0.583 |
| | ESM-2 + ESM-MSA-1b | 0.688 | 0.882 | 0.773 | 0.775 | 0.938 | 0.608 |
| | ESM-MSA-1b | 0.641 | 0.860 | 0.735 | 0.739 | 0.927 | 0.554 |
| | ESM-2 | 0.661 | 0.883 | 0.756 | 0.760 | 0.938 | 0.586 |
| | LMetalSite | N/A | N/A | N/A | N/A | N/A | N/A |
| $Fe^{2+}$ | PSSM | 0.842 | 0.345 | 0.489 | 0.536 | 0.672 | 0.297 |
| | ESM-2 + PSSM | 0.796 | 0.874 | 0.833 | 0.832 | 0.936 | 0.696 |
| | ESM-2 + ESM-MSA-1b | 0.749 | 0.846 | 0.795 | 0.794 | 0.922 | 0.636 |
| | ESM-MSA-1b | 0.746 | 0.714 | 0.730 | 0.727 | 0.856 | 0.535 |
| | ESM-2 | 0.868 | 0.849 | 0.858 | 0.857 | 0.924 | 0.739 |
| | LMetalSite | N/A | N/A | N/A | N/A | N/A | N/A |
| $Co^{2+}$ | PSSM | 0.000 | 0.000 | 0.000 | 0.000 | 0.500 | 0.012 |
| | ESM-2 + PSSM | 0.845 | 0.377 | 0.522 | 0.561 | 0.688 | 0.326 |

| | | | | | | | |
|---|---|---|---|---|---|---|---|
| | ESM-2 + ESM-MSA-1b | 0.935 | 0.346 | 0.505 | 0.566 | 0.673 | 0.331 |
| | ESM-MSA-1b | 1.000 | 0.277 | 0.434 | 0.524 | 0.638 | 0.286 |
| | ESM-2 | 0.918 | 0.464 | 0.616 | 0.650 | 0.732 | 0.432 |
| | LMetalSite | N/A | N/A | N/A | N/A | N/A | N/A |
| $Po_4^{3-}$ | PSSM | 0.000 | 0.000 | 0.000 | 0.000 | 0.500 | 0.017 |
| | ESM-2 + PSSM | 0.420 | 0.209 | 0.279 | 0.288 | 0.602 | 0.101 |
| | ESM-2 + ESM-MSA-1b | 0.404 | 0.170 | 0.239 | 0.254 | 0.583 | 0.083 |
| | ESM-MSA-1b | 0.341 | 0.119 | 0.177 | 0.194 | 0.558 | 0.056 |
| | ESM-2 | 0.369 | 0.197 | 0.256 | 0.260 | 0.595 | 0.086 |
| | LMetalSite | N/A | N/A | N/A | N/A | N/A | N/A |
| $So_4^{2-}$ | PSSM | 0.000 | 0.000 | 0.000 | 0.000 | 0.500 | 0.023 |
| | ESM-2 + PSSM | 0.513 | 0.154 | 0.237 | 0.272 | 0.575 | 0.099 |
| | ESM-2 + ESM-MSA-1b | 0.510 | 0.147 | 0.229 | 0.266 | 0.572 | 0.095 |
| | ESM-MSA-1b | 0.543 | 0.096 | 0.163 | 0.222 | 0.547 | 0.073 |
| | ESM-2 | 0.507 | 0.145 | 0.225 | 0.262 | 0.571 | 0.093 |
| | LMetalSite | N/A | N/A | N/A | N/A | N/A | N/A |

* Dimension of above embeddings: PSSM (L x 20); ESM-2 (L x 1280); ESM-2 + PSSM (L x 1300); ESM-MSA-1b (L x 768); ESM-2 + ESM-MSA-1b (2048)

**Table S4.** Benchmark on MIonSite benchmark set

| Metal | Method | Year | Sen (%) | Spe (%) | Acc (%) | MCC |
|---|---|---|---|---|---|---|
| $Zn^{2+}$ | MetalDetector | 2008 | 38.26 | 99.83 | 98.22 | 0.565 |
| | S-SITE | 2013 | 45.14 | 97.88 | 96.51 | 0.387 |
| | TargetS | 2013 | 41.70 | 99.80 | 98.29 | 0.588 |
| | COACH | 2013 | 32.39 | 99.37 | 97.62 | 0.422 |
| | IonSeq | 2016 | 70.04 | 92.56 | 91.97 | 0.347 |
| | MIB | 2016 | 40.12 | 99.07 | 97.53 | 0.451 |
| | IonCom | 2016 | 76.72 | 95.59 | 95.10 | 0.474 |
| | MIonSite | 2019 | 70.65 | 99.68 | 98.92 | 0.771 |
| | LMetalSite | 2022 | 78.72 | 99.81 | 99.37 | 0.836 |
| | GASS-Metal (1) | 2022 | 52.14 | 99.35 | 98.75 | 0.515 |
| | GASS-Metal (2) | 2022 | 65.19 | 99.49 | 98.96 | 0.647 |
| | GASS-Metal (3) | 2022 | 75.96 | 99.68 | 99.32 | 0.756 |
| | M-Ionic | Current | 76.64 | 99.50 | 99.04 | 0.760 |
| $Ca^{2+}$ | MetalDetector | 2008 | 0.62 | 99.86 | 98.03 | 0.016 |
| | S-SITE | 2013 | 10.17 | 99.51 | 97.87 | 0.159 |
| | TargetS | 2013 | 19.71 | 99.70 | 98.23 | 0.322 |
| | COACH | 2013 | 16.18 | 97.82 | 96.32 | 0.122 |
| | IonSeq | 2016 | 0.00 | 100 | 98.16 | N/A |
| | MIB | 2016 | 17.61 | 99.17 | 97.68 | 0.213 |
| | IonCom | 2016 | 28.01 | 99.47 | 98.16 | 0.365 |
| | MIonSite | 2019 | 42.53 | 99.71 | 98.66 | 0.552 |
| | LMetalSite | 2022 | 36.02 | 99.81 | 98.97 | 0.506 |
| | GASS-Metal (1) | 2022 | 9.38 | 98.84 | 97.60 | 0.081 |
| | GASS-Metal (2) | 2022 | 22.93 | 99.00 | 97.96 | 0.219 |
| | GASS-Metal (3) | 2022 | 29.05 | 99.09 | 98.15 | 0.281 |
| | M-Ionic | Current | 44.41 | 99.27 | 98.45 | 0.450 |
| $Mg^{2+}$ | MetalDetector | 2008 | 1.57 | 99.85 | 98.31 | 0.043 |
| | S-SITE | 2013 | 35.02 | 97.28 | 96.31 | 0.227 |
| | TargetS | 2013 | 15.16 | 99.84 | 98.52 | 0.298 |
| | COACH | 2013 | 21.12 | 97.60 | 96.41 | 0.144 |
| | IonSeq | 2016 | 0.00 | 100 | 98.44 | N/A |
| | MIB | 2016 | 22.21 | 99.33 | 98.12 | 0.268 |
| | IonCom | 2016 | 24.12 | 99.25 | 98.07 | 0.276 |
| | MIonSite | 2019 | 24.35 | 99.69 | 98.51 | 0.362 |
| | LMetalSite | 2022 | 24.88 | 99.87 | 99.03 | 0.410 |
| | GASS-Metal (1) | 2022 | 13.05 | 99.34 | 98.75 | 0.124 |

| | | | | | | |
|---|---|---|---|---|---|---|
| | GASS-Metal (2) | 2022 | 33.88 | 99.54 | 99.05 | 0.333 |
| | GASS-Metal (3) | 2022 | 55.74 | 99.69 | 99.36 | 0.550 |
| | M-Ionic | Current | 29.01 | 99.59 | 98.57 | 0.380 |
| $Mn^{2+}$ | MetalDetector | 2008 | 21.05 | 99.72 | 98.69 | 0.319 |
| | S-SITE | 2013 | 69.47 | 98.38 | 98.01 | 0.493 |
| | TargetS | 2013 | 28.42 | 99.75 | 98.82 | 0.408 |
| | COACH | 2013 | 27.01 | 99.82 | 98.87 | 0.420 |
| | IonSeq | 2016 | 2.11 | 99.94 | 98.67 | 0.081 |
| | MIB | 2016 | 47.75 | 99.52 | 98.84 | 0.514 |
| | IonCom | 2016 | 54.06 | 99.39 | 98.80 | 0.534 |
| | MIonSite | 2019 | 57.27 | 99.40 | 98.84 | 0.558 |
| | LMetalSite | 2022 | 67.78 | 99.81 | 99.41 | 0.740 |
| | GASS-Metal (1) | 2022 | 19.45 | 99.45 | 98.70 | 0.188 |
| | GASS-Metal (2) | 2022 | 43.75 | 99.60 | 99.10 | 0.433 |
| | GASS-Metal (3) | 2022 | 55.20 | 99.60 | 99.35 | 0.559 |
| | M-Ionic | Current | 58.46 | 99.80 | 99.29 | 0.680 |
| $Fe^{3+}$ | MetalDetector | 2008 | 28.57 | 99.76 | 99.24 | 0.360 |
| | S-SITE | 2013 | 90.48 | 98.78 | 98.71 | 0.560 |
| | TargetS | 2013 | 28.57 | 99.55 | 99.03 | 0.296 |
| | COACH | 2013 | 77.52 | 99.73 | 99.57 | 0.725 |
| | IonSeq | 2016 | 80.95 | 96.85 | 96.73 | 0.350 |
| | MIB | 2016 | 52.63 | 97.83 | 97.49 | 0.277 |
| | IonCom | 2016 | 77.12 | 99.80 | 99.64 | 0.756 |
| | MIonSite | 2019 | 82.90 | 99.75 | 99.63 | 0.765 |
| | LMetalSite | 2022 | 95.00 | 99.93 | 99.90 | 0.927 |
| | GASS-Metal (1) | 2022 | 73.40 | 99.97 | 99.85 | 0.733 |
| | GASS-Metal (2) | 2022 | 92.00 | 99.97 | 99.85 | 0.920 |
| | GASS-Metal (3) | 2022 | 92.00 | 99.97 | 99.85 | 0.920 |
| | M-Ionic | Current | 80.00 | 99.78 | 99.62 | 0.770 |
| $Cu^{2+}$ | MetalDetector | 2008 | 58.33 | 99.91 | 99.47 | 0.712 |
| | S-SITE | 2013 | 25.00 | 97.41 | 96.64 | 0.138 |
| | COACH | 2013 | 50.00 | 97.14 | 96.64 | 0.268 |
| | IonSeq | 2016 | 41.67 | 99.11 | 98.50 | 0.365 |
| | MIB | 2016 | 83.33 | 98.03 | 97.88 | 0.503 |
| | IonCom | 2016 | 50.00 | 99.55 | 99.03 | 0.517 |
| | MIonSite | 2019 | 48.02 | 99.63 | 99.08 | 0.523 |
| | LMetalSite | 2022 | 81.82 | 100 | 99.82 | 0.904 |
| | GASS-Metal (1) | 2022 | 67.00 | 99.93 | 99.85 | 0.666 |
| | GASS-Metal (2) | 2022 | 67.00 | 99.93 | 99.85 | 0.666 |

| | | | | | | |
|---|---|---|---|---|---|---|
| | GASS-Metal (3) | 2022 | 67.00 | 99.93 | 99.85 | 0.666 |
| | M-Ionic | Current | 54.55 | 100 | 99.56 | 0.740 |
| $Fe^{2+}$ | MetalDetector | 2008 | 33.33 | 99.86 | 99.04 | 0.496 |
| | S-SITE | 2013 | 66.67 | 95.54 | 95.18 | 0.309 |
| | COACH | 2013 | 66.67 | 98.05 | 97.66 | 0.437 |
| | IonSeq | 2016 | 97.80 | 99.28 | 99.26 | 0.782 |
| | MIB | 2016 | 100 | 99.44 | 99.45 | 0.830 |
| | IonCom | 2016 | 100 | 99.44 | 99.45 | 0.830 |
| | MIonSite | 2019 | 94.98 | 99.56 | 99.5 | 0.830 |
| | LMetalSite | 2022 | 100 | 100 | 100 | 1.000 |
| | GASS-Metal (1) | 2022 | 78.00 | 99.67 | 99.67 | 0.775 |
| | GASS-Metal (2) | 2022 | 89.00 | 99.67 | 99.67 | 0.887 |
| | GASS-Metal (3) | 2022 | 89.00 | 99.67 | 99.67 | 0.887 |
| | M-Ionic | Current | 100 | 100 | 100 | 1.000 |
| $Co^{2+}$ | MetalDetector | 2008 | 16.67 | 99.58 | 98.69 | 0.217 |
| | S-SITE | 2013 | 55.21 | 84.68 | 84.36 | 0.113 |
| | COACH | 2013 | 53.21 | 91.56 | 91.14 | 0.162 |
| | MIB | 2016 | 33.33 | 95.48 | 94.81 | 0.138 |
| | MIonSite | 2019 | 58.77 | 92.58 | 92.22 | 0.195 |
| | LMetalSite | 2022 | 22.22 | 100 | 99.17 | 0.469 |
| | GASS-Metal (1) | 2022 | 75.00 | 99.95 | 99.90 | 0.749 |
| | GASS-Metal (2) | 2022 | 75.00 | 99.95 | 99.90 | 0.749 |
| | GASS-Metal (3) | 2022 | 75.00 | 99.95 | 99.90 | 0.749 |
| | M-Ionic | Current | 0 | 100 | 98.60 | 0 |

*GASS-Metal (1) = top 10 results; GASS-Metal (2) = top 100 results; GASS-Metal (3) = all results from search

**Table S5.** Impact of structural features (DSSP) on metal-binding site prediction using the independent test set generated in this study (TestFold6)

| Ligand Type | Features | Precision | Recall | F1-score | MCC | AUROC | Average Precision |
|---|---|---|---|---|---|---|---|
| $Zn^{2+}$ | DSSP | 0.190 | 0.037 | 0.062 | 0.078 | 0.517 | 0.021 |
| | ESM2+DSSP | 0.678 | 0.798 | 0.733 | 0.732 | 0.896 | 0.544 |
| | LMetalSite | 0.856 | 0.780 | 0.816 | 0.814 | 0.889 | 0.671 |
| | ESM-2 | 0.739 | 0.869 | 0.799 | 0.798 | 0.932 | 0.645 |
| | ESM-MSA-1b | 0.712 | 0.825 | 0.764 | 0.763 | 0.910 | 0.590 |
| | ProtT5-XL | 0.600 | 0.200 | 0.300 | 0.333 | 0.597 | 0.150 |

| | | | | | | | |
|---|---|---|---|---|---|---|---|
| $Ca^{2+}$ | DSSP | 0.106 | 0.037 | 0.055 | 0.053 | 0.516 | 0.021 |
| | ESM2+DSSP | 0.392 | 0.430 | 0.410 | 0.400 | 0.709 | 0.179 |
| | LMetalSite | 0.815 | 0.364 | 0.504 | 0.540 | 0.681 | 0.308 |
| | ESM-2 | 0.527 | 0.549 | 0.538 | 0.529 | 0.770 | 0.297 |
| | ESM-MSA-1b | 0.267 | 0.435 | 0.331 | 0.326 | 0.707 | 0.126 |
| | ProtT5-XL | 0.000 | 0.000 | 0.000 | -0.003 | 0.499 | 0.005 |
| $Mg^{2+}$ | DSSP | 0.121 | 0.008 | 0.016 | 0.029 | 0.504 | 0.013 |
| | ESM2+DSSP | 0.432 | 0.337 | 0.379 | 0.375 | 0.666 | 0.154 |
| | LMetalSite | 0.733 | 0.281 | 0.406 | 0.450 | 0.640 | 0.214 |
| | ESM-2 | 0.482 | 0.434 | 0.457 | 0.451 | 0.714 | 0.216 |
| | ESM-MSA-1b | 0.434 | 0.327 | 0.373 | 0.370 | 0.661 | 0.150 |
| | ProtT5-XL | 1.000 | 0.400 | 0.571 | 0.629 | 0.700 | 0.410 |
| $Mn^{2+}$ | DSSP | 0.000 | 0.000 | 0.000 | 0.000 | 0.500 | 0.013 |
| | ESM2+DSSP | 0.763 | 0.582 | 0.660 | 0.663 | 0.790 | 0.450 |
| | LMetalSite | 0.830 | 0.715 | 0.768 | 0.768 | 0.856 | 0.598 |
| | ESM-2 | 0.781 | 0.726 | 0.753 | 0.750 | 0.862 | 0.571 |
| | ESM-MSA-1b | 0.729 | 0.595 | 0.655 | 0.654 | 0.796 | 0.439 |
| | ProtT5-XL | 0.500 | 0.500 | 0.500 | 0.485 | 0.742 | 0.265 |
| $Fe^{3+}$ | DSSP | 0.000 | 0.000 | 0.000 | 0.000 | 0.500 | 0.022 |
| | ESM2+DSSP | 0.776 | 0.612 | 0.684 | 0.683 | 0.804 | 0.483 |
| | LMetalSite | N/A | N/A | N/A | N/A | N/A | N/A |
| | ESM-2 | 0.755 | 0.862 | 0.805 | 0.802 | 0.928 | 0.653 |
| | ESM-MSA-1b | 0.748 | 0.384 | 0.507 | 0.529 | 0.690 | 0.300 |
| | ProtT5-XL | 0.667 | 0.333 | 0.444 | 0.465 | 0.665 | 0.233 |
| $Cu^{2+}$ | DSSP | 0.000 | 0.000 | 0.000 | 0.000 | 0.500 | 0.016 |
| | ESM2+DSSP | 0.509 | 0.711 | 0.593 | 0.594 | 0.850 | 0.366 |
| | LMetalSite | N/A | N/A | N/A | N/A | N/A | N/A |
| | ESM-2 | 0.709 | 0.835 | 0.767 | 0.766 | 0.915 | 0.595 |
| | ESM-MSA-1b | 0.581 | 0.790 | 0.670 | 0.672 | 0.890 | 0.462 |
| | ProtT5-XL | 0.600 | 0.750 | 0.667 | 0.667 | 0.872 | 0.453 |
| $Fe^{2+}$ | DSSP | 0.000 | 0.000 | 0.000 | 0.000 | 0.500 | 0.015 |
| | ESM2+DSSP | 0.727 | 0.755 | 0.741 | 0.737 | 0.875 | 0.553 |

| | | | | | | | |
|---|---|---|---|---|---|---|---|
| | LMetalSite | N/A | N/A | N/A | N/A | N/A | N/A |
| | ESM-2 | 0.798 | 0.691 | 0.741 | 0.739 | 0.844 | 0.556 |
| | ESM-MSA-1b | 0.765 | 0.798 | 0.781 | 0.778 | 0.897 | 0.614 |
| | ProtT5-XL | 0.889 | 1.000 | 0.941 | 0.937 | 0.993 | 0.889 |
| $Co^{2+}$ | DSSP | 0.000 | 0.000 | 0.000 | 0.000 | 0.500 | 0.016 |
| | ESM2+DSSP | 0.615 | 0.037 | 0.070 | 0.149 | 0.518 | 0.039 |
| | LMetalSite | N/A | N/A | N/A | N/A | N/A | N/A |
| | ESM-2 | 0.800 | 0.264 | 0.397 | 0.456 | 0.632 | 0.223 |
| | ESM-MSA-1b | 0.706 | 0.053 | 0.098 | 0.190 | 0.526 | 0.052 |
| | ProtT5-XL | 0.400 | 1.000 | 0.571 | 0.630 | 0.996 | 0.400 |
| $Po_4^{3-}$ | DSSP | 0.000 | 0.000 | 0.000 | 0.000 | 0.500 | 0.019 |
| | ESM2+DSSP | 0.410 | 0.244 | 0.306 | 0.306 | 0.618 | 0.114 |
| | LMetalSite | N/A | N/A | N/A | N/A | N/A | N/A |
| | ESM-2 | 0.433 | 0.347 | 0.385 | 0.377 | 0.669 | 0.162 |
| | ESM-MSA-1b | 0.396 | 0.186 | 0.253 | 0.262 | 0.590 | 0.089 |
| | ProtT5-XL | 0.000 | 0.000 | 0.000 | 0.000 | 0.500 | 0.005 |
| $So_4^{2-}$ | DSSP | 0.000 | 0.000 | 0.000 | 0.000 | 0.500 | 0.018 |
| | ESM2+DSSP | 0.392 | 0.124 | 0.189 | 0.213 | 0.560 | 0.064 |
| | LMetalSite | N/A | N/A | N/A | N/A | N/A | N/A |
| | ESM-2 | 0.528 | 0.254 | 0.343 | 0.358 | 0.625 | 0.148 |
| | ESM-MSA-1b | 0.520 | 0.098 | 0.166 | 0.221 | 0.548 | 0.068 |
| | ProtT5-XL | 0.000 | 0.000 | 0.000 | 0.000 | 0.500 | 0.020 |

**Table S6.** Validating that M-Ionic is trained on the residue level (on the embedding dimension, e.g. 1280 for ESM-2) and not on the protein level (on the length L of the protein) using the independent test set generated in this study (TestFold6)

| Ligand Type | Features | Precision | Recall | F1-score | MCC | AUROC | Average Precision |
|---|---|---|---|---|---|---|---|
| $Zn^{2+}$ | ESM-2 (Batch-1) | 0.806 | 0.724 | 0.763 | 0.760 | 0.861 | 0.588 |
| | ESM-2 (Batch-128) *(Base model)* | 0.765 | 0.776 | 0.771 | 0.767 | 0.886 | 0.598 |

| | | | | | | | |
|---|---|---|---|---|---|---|---|
| | ESM-2<br>(Scrambled Batch-128) | 0.744 | 0.769 | 0.756 | 0.753 | 0.883 | 0.576 |
| $Ca^{2+}$ | ESM-2<br>(Batch-1) | 0.542 | 0.357 | 0.431 | 0.432 | 0.676 | 0.205 |
| | ESM-2 (Batch-128)<br>*(Base model)* | 0.505 | 0.415 | 0.456 | 0.449 | 0.704 | 0.220 |
| | ESM-2<br>(Scrambled Batch-128) | 0.504 | 0.392 | 0.441 | 0.436 | 0.693 | 0.208 |
| $Mg^{2+}$ | ESM-2<br>(Batch-1) | 0.519 | 0.310 | 0.388 | 0.395 | 0.653 | 0.169 |
| | ESM-2 (Batch-128)<br>*(Base model)* | 0.484 | 0.329 | 0.392 | 0.393 | 0.662 | 0.167 |
| | ESM-2<br>(Scrambled Batch-128) | 0.457 | 0.296 | 0.359 | 0.361 | 0.646 | 0.144 |
| $Mn^{2+}$ | ESM-2<br>(Batch-1) | 0.852 | 0.490 | 0.622 | 0.642 | 0.744 | 0.424 |
| | ESM-2 (Batch-128)<br>*(Base model)* | 0.825 | 0.498 | 0.621 | 0.637 | 0.748 | 0.417 |
| | ESM-2<br>(Scrambled Batch-128) | 0.807 | 0.539 | 0.646 | 0.656 | 0.768 | 0.441 |
| $Fe^{3+}$ | ESM-2<br>(Batch-1) | 0.868 | 0.472 | 0.611 | 0.635 | 0.735 | 0.421 |
| | ESM-2 (Batch-128)<br>*(Base model)* | 0.807 | 0.447 | 0.575 | 0.594 | 0.722 | 0.373 |
| | ESM-2<br>(Scrambled Batch-128) | 0.713 | 0.524 | 0.604 | 0.604 | 0.760 | 0.384 |
| $Cu^{2+}$ | ESM-2<br>(Batch-1) | 0.754 | 0.690 | 0.720 | 0.717 | 0.843 | 0.525 |
| | ESM-2 (Batch-128)<br>*(Base model)* | 0.735 | 0.753 | 0.744 | 0.740 | 0.874 | 0.558 |
| | ESM-2<br>(Scrambled Batch-128) | 0.691 | 0.761 | 0.724 | 0.721 | 0.878 | 0.530 |

| | | | | | | | |
|---|---|---|---|---|---|---|---|
| $Fe^{2+}$ | ESM-2 (Batch-1) | 0.822 | 0.686 | 0.748 | 0.748 | 0.842 | 0.568 |
| | ESM-2 (Batch-128) *(Base model)* | 0.802 | 0.713 | 0.755 | 0.753 | 0.855 | 0.576 |
| | ESM-2 (Scrambled Batch-128) | 0.767 | 0.702 | 0.733 | 0.730 | 0.849 | 0.543 |
| $Co^{2+}$ | ESM-2 (Batch-1) | 0.733 | 0.048 | 0.091 | 0.186 | 0.524 | 0.051 |
| | ESM-2 (Batch-128) *(Base model)* | 0.591 | 0.057 | 0.104 | 0.181 | 0.528 | 0.049 |
| | ESM-2 (Scrambled Batch-128) | 0.739 | 0.075 | 0.136 | 0.232 | 0.537 | 0.070 |
| $Po_4^{3-}$ | ESM-2 (Batch-1) | 0.362 | 0.108 | 0.166 | 0.190 | 0.552 | 0.056 |
| | ESM-2 (Batch-128) *(Base model)* | 0.442 | 0.157 | 0.232 | 0.255 | 0.577 | 0.085 |
| | ESM-2 (Scrambled Batch-128) | 0.373 | 0.159 | 0.223 | 0.234 | 0.577 | 0.075 |
| $So_4^{2-}$ | ESM-2 (Batch-1) | 0.513 | 0.076 | 0.132 | 0.192 | 0.537 | 0.056 |
| | ESM-2 (Batch-128) *(Base model)* | 0.385 | 0.076 | 0.127 | 0.164 | 0.537 | 0.046 |
| | ESM-2 (Scrambled Batch-128) | 0.489 | 0.087 | 0.148 | 0.201 | 0.543 | 0.060 |

**Table S7:** Analysis of M-Ionic performance for each ion for each taxon

| Ligand Type | Taxonomy | Precision | Recall | F1-score | MCC | AUROC |
|---|---|---|---|---|---|---|
| $Zn^{2+}$ | Archaea | 0.734 | 0.972 | 0.814 | 0.827 | 0.978 |
| | Bacteria | 0.664 | 0.832 | 0.715 | 0.725 | 0.909 |
| | Eukaryota | 0.829 | 0.942 | 0.870 | 0.871 | 0.964 |
| | Viruses | 0.623 | 0.815 | 0.687 | 0.695 | 0.901 |
| $Ca^{2+}$ | Archaea | 0.383 | 0.708 | 0.428 | 0.447 | 0.817 |
| | Bacteria | 0.335 | 0.433 | 0.358 | 0.361 | 0.709 |

| | | | | | | |
|---|---|---|---|---|---|---|
| | Eukaryota | 0.520 | 0.696 | 0.560 | 0.565 | 0.829 |
| | Viruses | 0.372 | 0.588 | 0.436 | 0.445 | 0.782 |
| $Mg^{2+}$ | Archaea | 0.321 | 0.591 | 0.380 | 0.404 | 0.784 |
| | Bacteria | 0.301 | 0.513 | 0.356 | 0.371 | 0.749 |
| | Eukaryota | 0.421 | 0.665 | 0.487 | 0.509 | 0.828 |
| | Viruses | 0.416 | 0.594 | 0.443 | 0.469 | 0.795 |
| $Mn^{2+}$ | Archaea | 0.783 | 1.000 | 0.878 | 0.883 | 0.998 |
| | Bacteria | 0.680 | 0.633 | 0.625 | 0.638 | 0.816 |
| | Eukaryota | 0.634 | 0.597 | 0.581 | 0.594 | 0.797 |
| | Viruses | 0.665 | 0.867 | 0.729 | 0.740 | 0.927 |
| $Fe^{3+}$ | Archaea | 0.690 | 0.521 | 0.551 | 0.571 | 0.760 |
| | Bacteria | 0.591 | 0.576 | 0.550 | 0.564 | 0.787 |
| | Eukaryota | 0.691 | 0.665 | 0.651 | 0.659 | 0.831 |
| $Cu^{2+}$ | Bacteria | 0.648 | 0.859 | 0.721 | 0.731 | 0.925 |
| | Eukaryota | 0.625 | 0.799 | 0.679 | 0.692 | 0.896 |
| $Fe^{2+}$ | Bacteria | 0.813 | 0.826 | 0.768 | 0.790 | 0.911 |
| | Eukaryota | 0.920 | 0.783 | 0.815 | 0.831 | 0.891 |
| $Co^{2+}$ | Bacteria | 0.445 | 0.292 | 0.338 | 0.351 | 0.646 |
| | Eukaryota | 0.389 | 0.468 | 0.421 | 0.424 | 0.733 |
| $Po_4^{3-}$ | Archaea | 0.399 | 0.167 | 0.229 | 0.246 | 0.581 |
| | Bacteria | 0.224 | 0.190 | 0.187 | 0.187 | 0.591 |
| | Eukaryota | 0.246 | 0.196 | 0.203 | 0.205 | 0.595 |
| | Viruses | 0.150 | 0.117 | 0.124 | 0.120 | 0.555 |
| $So_4^{2-}$ | Archaea | 0.204 | 0.122 | 0.131 | 0.134 | 0.552 |
| | Bacteria | 0.165 | 0.109 | 0.119 | 0.122 | 0.553 |
| | Eukaryota | 0.157 | 0.115 | 0.113 | 0.118 | 0.556 |
| | Viruses | 0.063 | 0.025 | 0.036 | 0.039 | 0.512 |

**Table S8:** Analysis of M-Ionic performance for each ion for DNA-binding proteins

| Ligand Type | Type | Precision | Recall | F1-score | MCC | AUROC | Average Precision |
|---|---|---|---|---|---|---|---|
| $Zn^{2+}$ (p-value = 0.0048) | non-binding | 0.780 | 0.911 | 0.824 | 0.828 | 0.949 | 0.758 |
| | DNA-binding | 0.867 | 0.980 | 0.913 | 0.913 | 0.985 | 0.863 |
| $Ca^{2+}$ (p-value = 0.7264) | non-binding | 0.451 | 0.606 | 0.487 | 0.492 | 0.788 | 0.390 |
| | DNA-binding | 0.500 | 0.333 | 0.400 | 0.403 | 0.665 | 0.342 |
| $Mg^{2+}$ (p-value = 0.6854) | non-binding | 0.371 | 0.599 | 0.430 | 0.450 | 0.794 | 0.332 |
| | DNA-binding | 0.307 | 0.551 | 0.370 | 0.395 | 0.773 | 0.304 |

| | | | | | | | |
|---|---|---|---|---|---|---|---|
| $Mn^{2+}$ (p-value = 1) | non-binding | 0.660 | 0.647 | 0.620 | 0.633 | 0.822 | 0.540 |
| | DNA-binding | 0.800 | 0.800 | 0.800 | 0.793 | 0.897 | 0.647 |
| $Fe^{3+}$ (p-value = N/A) | non-binding | 0.629 | 0.599 | 0.581 | 0.593 | 0.798 | 0.519 |
| | DNA-binding | - | - | - | - | - | - |
| $Cu^{2+}$ (p-value = N/A) | non-binding | 0.638 | 0.834 | 0.704 | 0.715 | 0.913 | 0.605 |
| | DNA-binding | - | - | - | - | - | - |
| $Fe^{2+}$ (p-value = 0.3420) | non-binding | 0.862 | 0.801 | 0.786 | 0.806 | 0.899 | 0.712 |
| | DNA-binding | 1.000 | 1.000 | 1.000 | 1.000 | 1.000 | 1.000 |
| $Co^{2+}$ (p-value = 0.3914) | non-binding | 0.424 | 0.395 | 0.391 | 0.399 | 0.697 | 0.346 |
| | DNA-binding | 0.000 | 0.000 | 0.000 | 0.000 | 0.500 | 0.010 |
| $Po_4^{3-}$ (p-value = 0.6437) | non-binding | 0.242 | 0.191 | 0.197 | 0.198 | 0.592 | 0.147 |
| | DNA-binding | 0.077 | 0.333 | 0.125 | 0.148 | 0.657 | 0.100 |
| $So_4^{2-}$ (p-value = 0.0878) | non-binding | 0.159 | 0.108 | 0.113 | 0.117 | 0.552 | 0.104 |
| | DNA-binding | 0.000 | 0.000 | 0.000 | -0.019 | 0.497 | 0.070 |

**Table S9:** Analysis of M-Ionic performance for each ion for transmembrane against non-membrane proteins

| Ligand Type | Type | Precision | Recall | F1-score | MCC | AUROC | Average Precision |
|---|---|---|---|---|---|---|---|
| $Zn^{2+}$ (p-value = 0.0054) | Transmembrane | 0.791 | 0.919 | 0.836 | 0.839 | 0.953 | 0.771 |
| | Non-membrane | 0.692 | 0.851 | 0.741 | 0.750 | 0.920 | 0.664 |
| $Ca^{2+}$ (p-value = 0.2037) | Transmembrane | 0.436 | 0.605 | 0.478 | 0.483 | 0.786 | 0.381 |
| | Non-membrane | 0.510 | 0.605 | 0.521 | 0.525 | 0.791 | 0.425 |
| $Mg^{2+}$ (p-value = 0.6428) | Transmembrane | 0.369 | 0.595 | 0.428 | 0.447 | 0.791 | 0.331 |
| | Non-membrane | 0.366 | 0.629 | 0.436 | 0.462 | 0.811 | 0.330 |
| $Mn^{2+}$ (p-value = 0.2683) | Transmembrane | 0.666 | 0.660 | 0.630 | 0.643 | 0.828 | 0.551 |
| | Non-membrane | 0.597 | 0.500 | 0.513 | 0.528 | 0.749 | 0.411 |
| $Fe^{3+}$ (p-value = 0.5236) | Transmembrane | 0.624 | 0.589 | 0.572 | 0.584 | 0.793 | 0.510 |
| | Non-membrane | 0.656 | 0.656 | 0.632 | 0.643 | 0.828 | 0.567 |
| $Cu^{2+}$ (p-value = 0.0369) | Transmembrane | 0.617 | 0.845 | 0.695 | 0.707 | 0.918 | 0.585 |
| | Non-membrane | 0.726 | 0.789 | 0.741 | 0.747 | 0.893 | 0.691 |
| $Fe^{2+}$ (p-value = 0.3459) | Transmembrane | 0.884 | 0.823 | 0.807 | 0.827 | 0.910 | 0.735 |
| | Non-membrane | 0.667 | 0.625 | 0.617 | 0.630 | 0.81Average Precision2 | 0.544 |
| $Co^{2+}$ (p-value = 0.11805) | Transmembrane | 0.442 | 0.413 | 0.408 | 0.416 | 0.706 | 0.361 |
| | Non-membrane | 0.000 | 0.000 | 0.000 | 0.000 | 0.500 | 0.010 |
| $Po_4^{3-}$ (p-value = | Transmembrane | 0.243 | 0.197 | 0.201 | 0.202 | 0.595 | 0.151 |

| | | | | | | | |
|---|---|---|---|---|---|---|---|
| 0.3561) | Non-membrane | 0.190 | 0.085 | 0.101 | 0.109 | 0.541 | 0.066 |
| $So_4^{2-}$ (p-value = 0.3023) | Transmembrane | 0.158 | 0.111 | 0.115 | 0.118 | 0.553 | 0.106 |
| | Non-membrane | 0.150 | 0.053 | 0.079 | 0.089 | 0.527 | 0.077 |

### 2. Figures

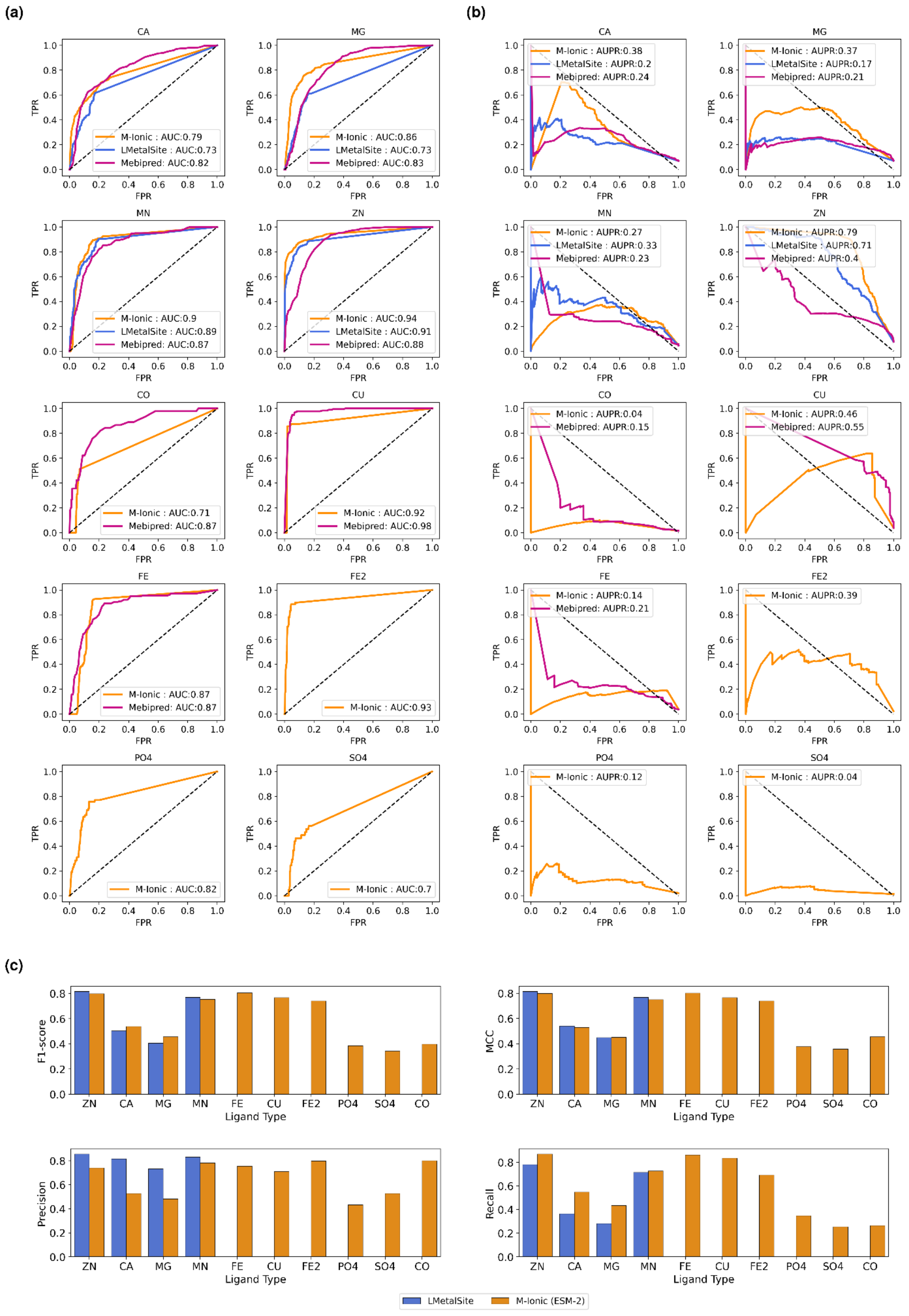

#### Figure. S1. Protein- and residue-level comparison on homology reduced independent test set (TestFold6)

(a) Protein-level: Comparison of ROC curves for the performance of each ion type for M-Ionic (this study) trained on ESM-2 embeddings, LMetalSite (Yuan et al., 2022) and mebi-pred (Aptekmann et al., 2022) on homology reduced independent test set (TestFold6) and the negative binding test set (b) Protein-level: Comparison of Precision-Recall curves for the performance of each ion type for each of the methods (c) Residue-level: F1, MCC, Precision and Recall scores of performance of M-Ionic (trained on ESM-2 and ESM-MSA-1b embeddings) and LMetalSite (Yuan et al., 2022) on homology reduced independent test set (TestFold6).

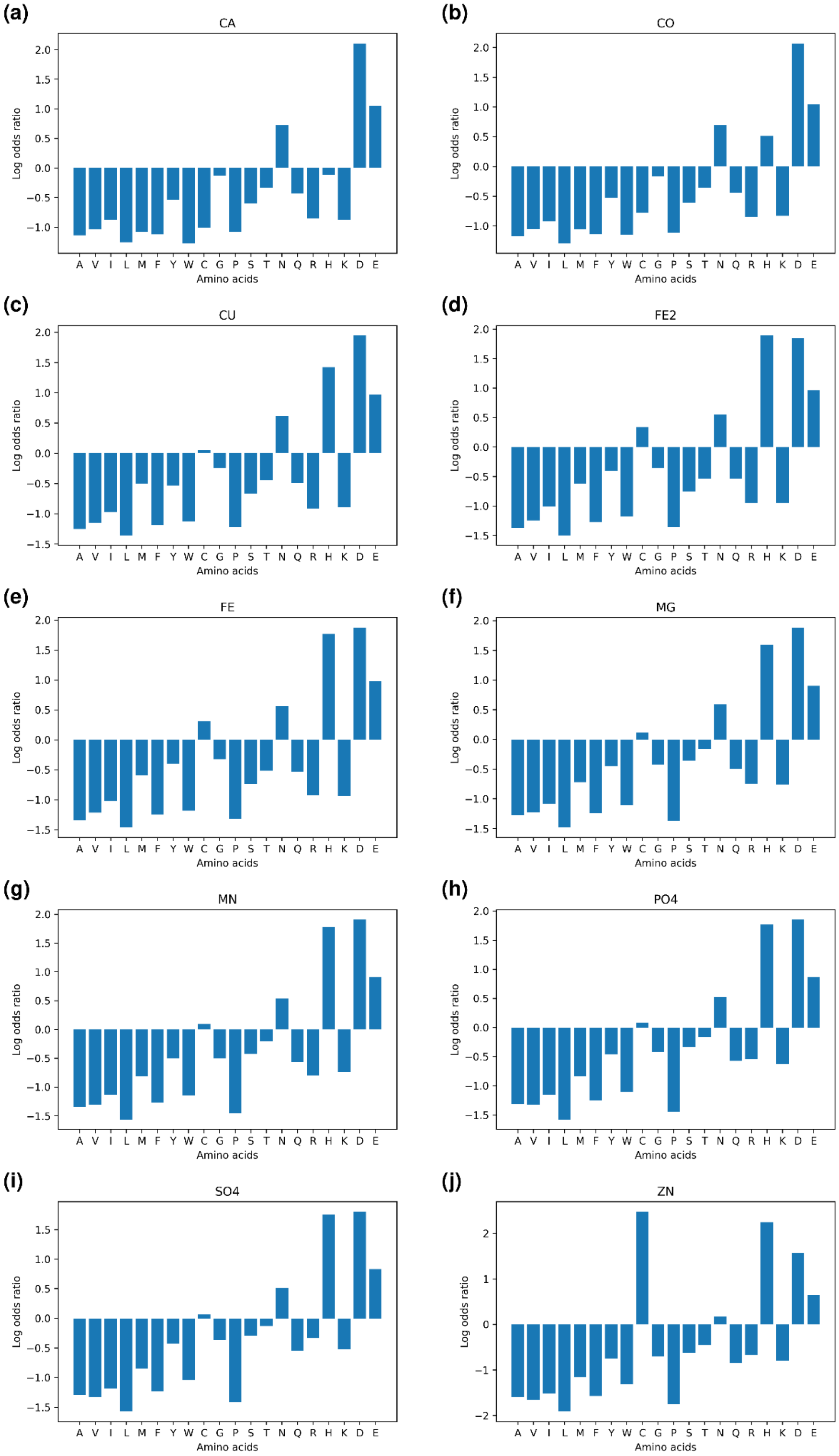

**Figure. S2.** Log odds ratio showing the binding propensity of amino acids with respect to each ion group. Positive log odds signify that certain amino acids are more likely to bind to that metal group, whereas a negative log odds ratio shows a non-preferential binding.

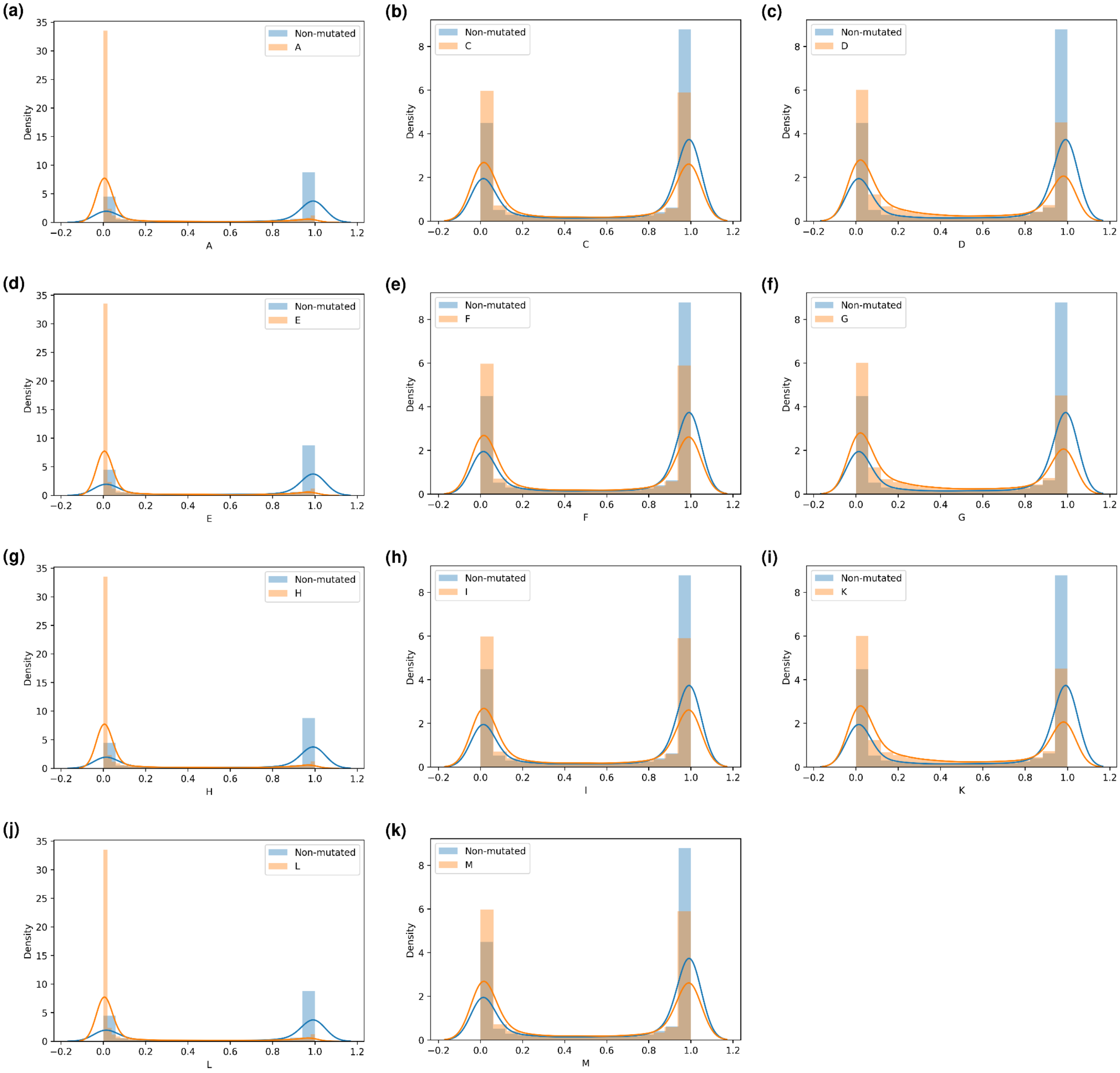

(a)
Non-mutated
A
Density
A
(b)
Non-mutated
C
Density
C
(c)
Non-mutated
D
Density
D
(d)
Non-mutated
E
Density
E
(e)
Non-mutated
F
Density
F
(f)
Non-mutated
G
Density
G
(g)
Non-mutated
H
Density
H
(h)
Non-mutated
I
Density
I
(i)
Non-mutated
K
Density
K
(j)
Non-mutated
L
Density
L
(k)
Non-mutated
M
Density
M

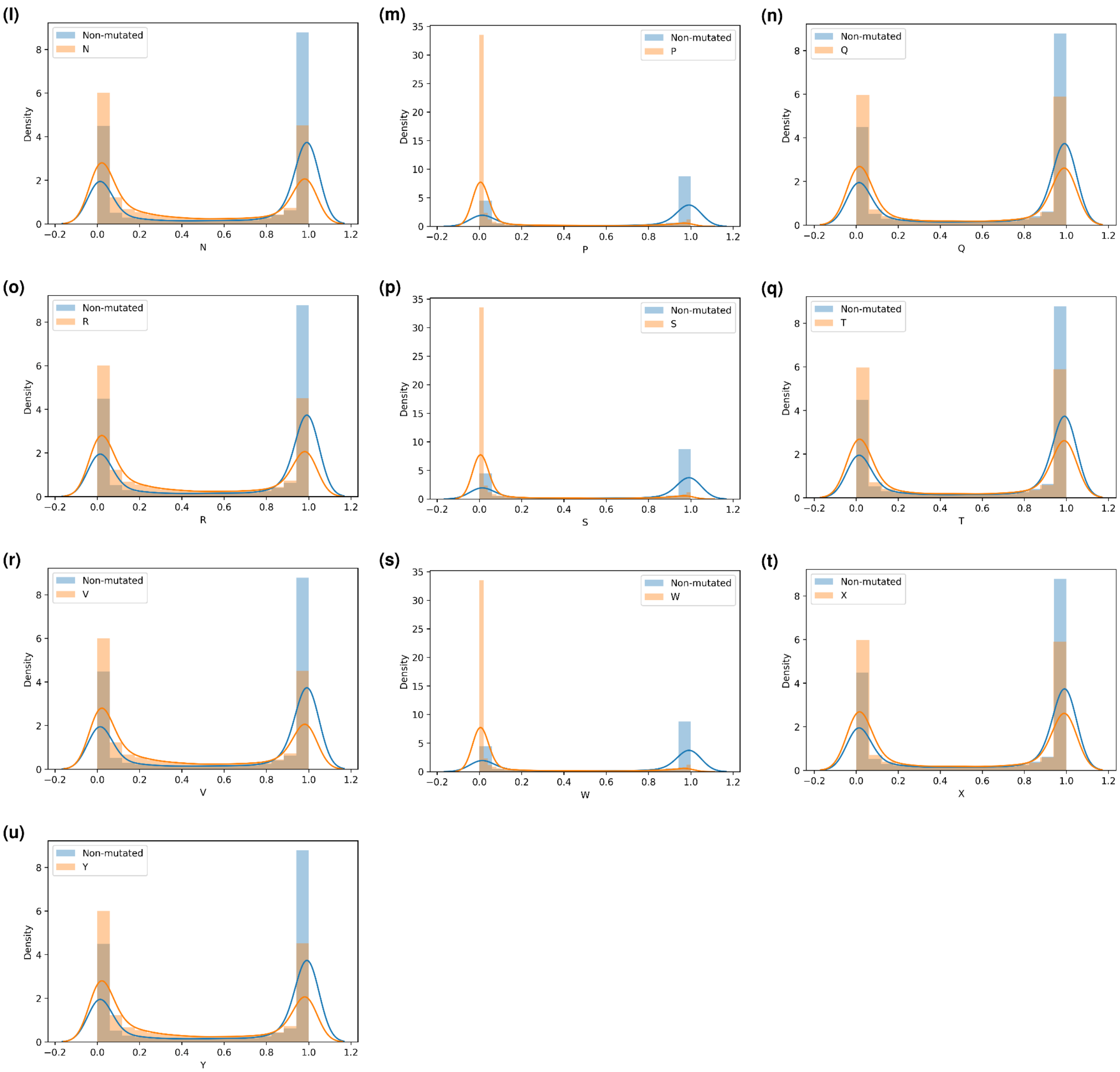

**Figure. S3.** Probabilities distributions (output from M-Ionic) showing the effect of mutating metal-binding residues to another residue compared to the non-mutated (original) sequence.
